## Supplementary material for "Foxtrot migration and dynamic over-wintering range of an arctic raptor"

* *Corresponding author*

### Supplementary Tables

**Table S1**. The relationship between latitude/longitude and the day of the year during quick and slow phases of migration. The likelihood ratio test compares two candidate models: with ("~doy") and without the day of the year ("~1") as a fixed factor. For the fixed effects, see Table S2. For the random effects, see Table S3.

| Migration | Model | df | AIC | BIC | logLik | dev | Chisq | p-value |
|---|---|---|---|---|---|---|---|---|
| Quick: spring (latitude) | ~ 1 | 4 | 4290 | 4309 | -2141 | 4282 | | |
| | ~ doy | 5 | 3612 | 3635 | -1801 | 3602 | 680 | <0.001 |
| Quick: fall (latitude) | ~ 1 | 4 | 4816 | 4834 | -2404 | 4808 | | |
| | ~ doy | 5 | 4277 | 4300 | -2133 | 4267 | 541 | <0.001 |
| Slow: 1st phase (latitude) | ~ 1 | 4 | 14916 | 14940 | -7454 | 14908 | | |
| | ~ doy | 5 | 13690 | 13721 | -6840 | 13680 | 1228 | <0.001 |
| Slow: 1st phase (longitude) | ~ 1 | 4 | 19746 | 19771 | -9869 | 19738 | | |
| | ~ doy | 5 | 18929 | 18960 | -9459 | 18919 | 820 | <0.001 |
| Slow: 2nd phase (latitude) | ~ 1 | 4 | 13561 | 13585 | -6777 | 13553 | | |
| | ~ doy | 5 | 11726 | 11756 | -5858 | 11716 | 1837 | <0.001 |
| Slow: 2nd phase (longitude) | ~ 1 | 4 | 19250 | 19275 | -9621 | 19242 | | |
| | ~ doy | 5 | 17482 | 17512 | -8736 | 17472 | 1770 | <0.001 |

**Table S2.** The relationship between latitude/longitude and the day of the year during quick and slow phases of migration. Linear mixed-effect model, fixed effects. The response variable – latitude/longitude. Fixed effect – the day of the year ('doy'). Random effects – individuals and year. For the random effects, see Table S3. For the likelihood ratio test results to compare candidate models, see Table S1.

| Migration | | Estimate | Std.Error | t value | p-value |
|---|---|---|---|---|---|
| Quick: spring (latitude) | Intercept | 2.945 | 1.928 | 1.53 | |
| | doy | 0.477 | 0.014 | 34.56 | <0.001 |
| Quick: fall (latitude) | Intercept | 201.875 | 4.872 | 41.44 | |
| | doy | -0.503 | 0.016 | -30.47 | <0.001 |
| Slow: 1st phase (latitude) | Intercept | 55.500 | 0.678 | 81.90 | |
| | doy | -0.026 | 0.001 | -38.55 | <0.001 |
| Slow: 2nd phase (latitude) | Intercept | 42.785 | 0.482 | 88.73 | |
| | doy | 0.046 | 0.001 | 50.03 | <0.001 |
| Slow: 1st phase (longitude) | Intercept | 46.186 | 1.764 | 26.18 | |
| | doy | -0.045 | 0.001 | -30.42 | <0.001 |
| Slow: 2nd phase (longitude) | Intercept | 18.566 | 1.518 | 12.23 | |
| | doy | 0.113 | 0.002 | 48.84 | <0.001 |

**Table S3.** The relationship between latitude/longitude and the day of the year during quick and slow phases of migration. Linear mixed-effect model, random effects. The response variable – latitude/longitude. Fixed effect – the day of the year. Random effects – individuals ('bird') and year ('year'). For the fixed effects, see Table S2. For the likelihood ratio test results to compare candidate models, see Table S1.

| Migration | Group | Variance | Std. Dev. |
|---|---|---|---|
| Quick: spring (latitude) | bird | 7.998 | 2.828 |
| | year | 1.579 | 1.256 |
| | residual | 7.054 | 2.656 |
| Quick: fall (latitude) | bird | 25.858 | 5.085 |
| | year | 12.766 | 3.573 |
| | residual | 8.849 | 2.975 |
| Slow: 1st phase (latitude) | bird | 6.812 | 2.610 |
| | year | 1.377 | 1.173 |
| | residual | 2.968 | 1.723 |
| Slow: 1st phase (longitude) | bird | 34.222 | 5.850 |
| | year | 14.999 | 3.873 |
| | residual | 13.547 | 3.681 |
| Slow: 2nd phase (latitude) | bird | 1.921 | 1.386 |
| | year | 0.959 | 0.979 |
| | residual | 2.424 | 1.557 |
| Slow: 2nd phase (longitude) | bird | 19.047 | 4.364 |
| | year | 10.699 | 3.271 |
| | residual | 15.344 | 3.917 |

**Table S4.** The difference between the distance of slow and quick migrations. Linear mixed-effect model, post-hoc results. The response variable – distance (km). Fixed effect – the type of migrations. Results of the post hoc comparison. For the plot see Figure 1c.

| Migration | difference | SE | df | t ratio | p-value |
|---|---|---|---|---|---|
| Quick (spring) – Quick (fall) | 210 | 81 | 171 | 2,58 | 0,053 |
| Quick (spring) – Slow (first) | 528 | 87 | 164 | 6,04 | <0.001 |
| Quick (spring) – Slow (second) | 502 | 87 | 164 | 5,74 | <0.001 |
| Quick (fall) – Slow (first) | 318 | 80 | 164 | 4,00 | <0.001 |
| Quick (fall) – Slow (second) | 292 | 80 | 164 | 3,68 | 0,002 |
| Slow (first) – Slow (second) | -26 | 85 | 154 | -0,30 | 0,990 |

**Table S5.** The difference between the duration of slow and quick migrations. Linear mixed-effect model, posthoc results. The response variable – duration (days). Fixed effect – the type of migrations. Results of the post hoc comparison. For the plot see Figure 1c.

| Migration | difference | SE | df | t ratio | p-value |
|---|---|---|---|---|---|
| Quick (spring) – Quick (fall) | 4 | 6 | 160 | 0,67 | 0,910 |
| Quick (spring) – Slow (first) | -103 | 6 | 153 | -16,99 | <0.001 |
| Quick (spring) – Slow (second) | -49 | 7 | 151 | -6,90 | <0.001 |
| Quick (fall) – Slow (first) | -107 | 6 | 154 | -19,36 | <0.001 |
| Quick (fall) – Slow (second) | -52 | 7 | 159 | -7,90 | <0.001 |
| Slow (first) – Slow (second) | 54 | 7 | 150 | 7,75 | <0.001 |

**Table S6.** The difference between the speed of slow and quick migrations. Linear mixed-effect model, posthoc results. The response variable – speed (km/day). Fixed effect – the type of migrations. Results of the post hoc comparison. For the plot, see Figure 1c.

| Migration | difference | SE | df | t ratio | p-value |
|---|---|---|---|---|---|
| Quick (spring) – Quick (fall) | -7 | 7 | 149 | -1.02 | 0.737 |
| Quick (spring) – Slow (first) | 92 | 7 | 143 | 12.05 | <0.001 |
| Quick (spring) – Slow (second) | 83 | 9 | 138 | 9.50 | <0.001 |
| Quick (fall) – Slow (first) | 99 | 7 | 143 | 14.33 | <0.001 |
| Quick (fall) – Slow (second) | 90 | 8 | 141 | 10.85 | <0.001 |
| Slow (first) – Slow (second) | -9 | 9 | 136 | -1.04 | 0.727 |

**Table S7.** The difference between the direction of the spring and the second phase of the winter migration and between the autumn and the first phase of the winter migration (two models). Linear mixed-effect models, posthoc results. The response variable in both models – direction (deg). Fixed effect – the type of migrations. Results of the post hoc comparison. For the plot, see Figure 1c.

| Migration | difference | SE | df | t ratio | p-value |
|---|---|---|---|---|---|
| Quick (spring) – Slow (second) | -50 | 3 | 70 | -16.08 | <0.001 |
| Quick (fall) – Slow (first) | -54 | 3 | 82 | -15,73 | <0.001 |

**Table S8.** The relationship between migration distance and the sex of the birds. The likelihood ratio test compares two candidate models: with ("~sex") and without the sex ("~1") as a fixed factor.

| Model | df | AIC | BIC | logLik | dev | Chisq | p-value |
|---|---|---|---|---|---|---|---|
| ~ 1 | 3 | 7673.8 | 7684.6 | -3833.9 | 7667.8 | | |
| ~ sex | 45 | 7674.7 | 7689.0 | -3833.3 | 7666.7 | 1.175 | 0.278 |

**Table S9.** The difference between vegetation land cover types crossed during quick (fall and spring) and slow (winter) migrations. General linear mixed-effect models. Results are given on the logit scale. For the plot, see Figure 2.

| Type | Migration | Estimate | SE | z value | p-value |
|---|---|---|---|---|---|
| Grassland | (Intercept) | -1.55 | 0.23 | -6.66 | |
| | Slow migration | 2.27 | 0.10 | 23.91 | <0.001 |
| Croplands | (Intercept) | -7.09 | 0.76 | -9.37 | |
| | Slow migration | 4.59 | 0.47 | 9.80 | <0.001 |
| Forests | (Intercept) | -0.22 | 0.15 | -1.49 | |
| | Slow migration | -3.04 | 0.11 | -28.49 | <0.001 |
| Urban | (Intercept) | -7.36 | 0.94 | -7.84 | |
| | Slow migration | 1.08 | 0.65 | 1.64 | 0.1 |

**Table S10.** The difference between snow cover conditions in the real situation ('Real') and two and two hypothetical situations – if birds spend winter in the place where they have arrived after fall migration ('Hyp stay') and if birds fly directly to the Southwest and stay there all winter ('Hyp SW'). General linear mixed-effect models. Results are given on the logit scale. Results of the post hoc comparison. For the plot, see Figure 3b.

| Month | Migration | Estimate | SE | z value | p-value |
|---|---|---|---|---|---|
| October | Hyp SW – Hyp stay | -1.08 | 0.20 | -5.45 | <0.001 |
| | Hyp SW – Real | -1.13 | 0.22 | -5.13 | <0.001 |
| | Hyp stay – Real | -0.05 | 0.18 | -0.30 | 0.952 |
| November | Hyp SW – Hyp stay | -0.75 | 0.09 | -8.38 | <0.001 |
| | Hyp SW – Real | -0.66 | 0.11 | -6.15 | <0.001 |
| | Hyp stay – Real | 0.10 | 0.10 | 0.97 | 0.595 |
| December | Hyp SW – Hyp stay | -1.18 | 0.07 | -16.28 | <0.001 |
| | Hyp SW – Real | -0.35 | 0.08 | -4.25 | <0.001 |
| | Hyp stay – Real | 0.83 | 0.09 | 9.56 | <0.001 |
| January | Hyp SW – Hyp stay | -2.51 | 0.15 | -16.63 | <0.001 |
| | Hyp SW – Real | 0.13 | 0.11 | 1.13 | 0.497 |
| | Hyp stay – Real | 2.63 | 0.16 | 16.07 | <0.001 |
| February | Hyp SW – Hyp stay | -4.88 | 0.45 | -10.80 | <0.001 |
| | Hyp SW – Real | -0.38 | 0.11 | -3.48 | 0.0015 |
| | Hyp stay – Real | 4.50 | 0.46 | 9.86 | <0.001 |
| March | Hyp SW – Hyp stay | -4.87 | 0.14 | -34.14 | <0.001 |
| | Hyp SW – Real | -1.81 | 0.11 | -16.94 | <0.001 |
| | Hyp stay – Real | 3.06 | 0.14 | 22.43 | <0.001 |
| April | Hyp SW – Hyp stay | -2.45 | 0.14 | -17.98 | <0.001 |
| | Hyp SW – Real | -2.47 | 0.15 | -1706 | <0.001 |
| | Hyp stay – Real | -0.03 | 0.09 | -0.29 | 0.955 |

### Supplementary Figures

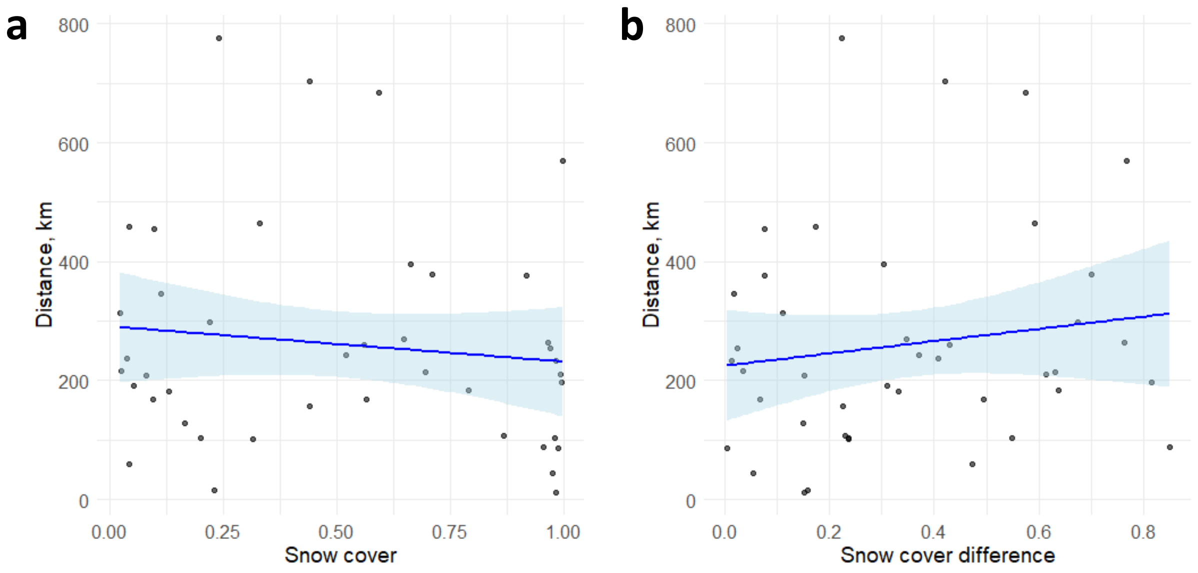


**Figure S1.** The distance between two consecutive monthly MCPs during the overwintering period did not depend on a) the snow cover extent (p=0.45) or b) the difference in snow cover (p=0.36).

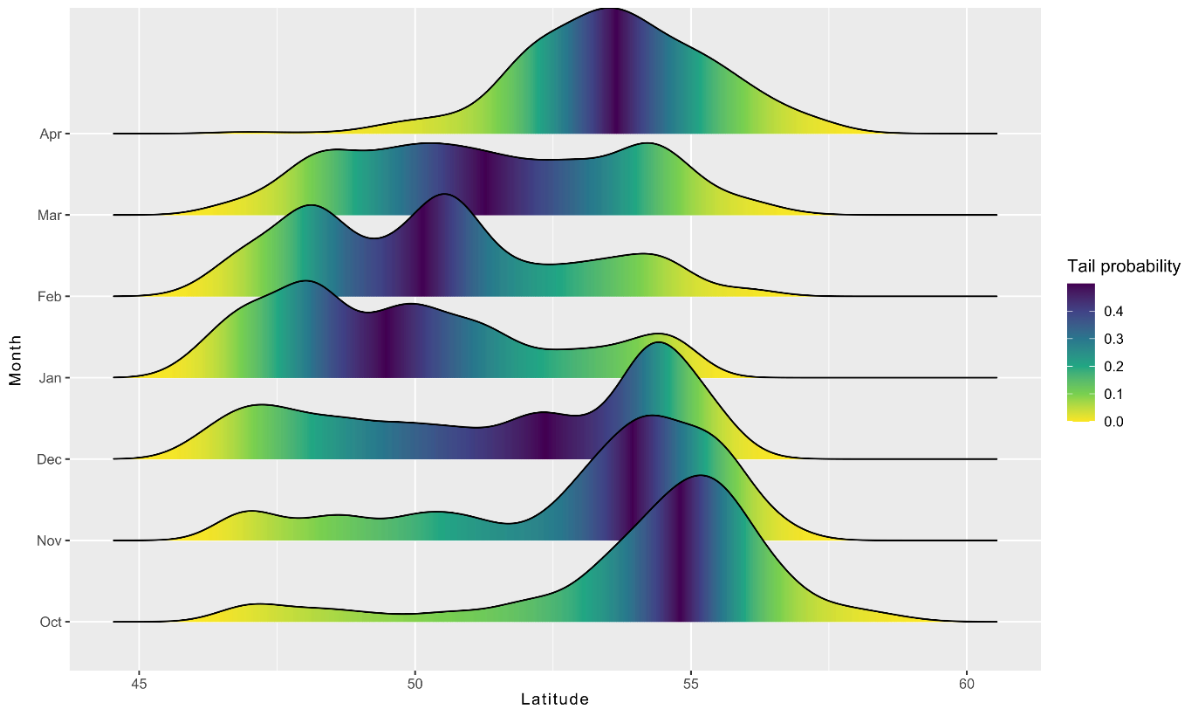


**Figure S2.** The distribution of latitudes of Rough-legged buzzards in different months.
